# Variation in PU.1 binding and chromatin looping at neutrophil enhancers influences autoimmune disease susceptibility

**DOI:** 10.1101/620260

**Authors:** Stephen Watt, Louella Vasquez, Klaudia Walter, Alice L. Mann, Kousik Kundu, Lu Chen, Ying Yan, Simone Ecker, Frances Burden, Samantha Farrow, Ben Farr, Valentina Iotchkova, Heather Elding, Daniel Mead, Manuel Tardaguila, Hannes Ponstingl, David Richardson, Avik Datta, Paul Flicek, Laura Clarke, Kate Downes, Tomi Pastinen, Peter Fraser, Mattia Frontini, Biola-Maria Javierre, Mikhail Spivakov, Nicole Soranzo

## Abstract

Neutrophils play fundamental roles in innate inflammatory response, shape adaptive immunity^1^, and have been identified as a potentially causal cell type underpinning genetic associations with immune system traits and diseases^2,3^ The majority of these variants are non-coding and the underlying mechanisms are not fully understood. Here, we profiled the binding of one of the principal myeloid transcriptional regulators, PU.1, in primary neutrophils across nearly a hundred volunteers, and elucidate the coordinated genetic effects of PU.1 binding variation, local chromatin state, promoter-enhancer interactions and gene expression. We show that PU.1 binding and the associated chain of molecular changes underlie genetically-driven differences in cell count and autoimmune disease susceptibility. Our results advance interpretation for genetic loci associated with neutrophil biology and immune disease.

## Results

Non-coding DNA sequence variation affects chromatin state and gene expression within human populations, and accounts for the majority of complex genetic traits and disease associations^4–7^. The commonly accepted model of genetic control of transcriptional activity postulates that genetic variation modifies the DNA recognition sequences of specific transcription factors (TFs), thus altering their ability to bind to DNA at a specific locus^8–15^. PU.1 (encoded by Spi1) is a key TF regulating myeloid development^16–18^, and its deficiency has profound effects on neutrophil maturation and function^19,20^. To study genetically determined variation in PU.1 recruitment to DNA, we used chromatin immunoprecipitation sequencing (ChIP-seq) to profile PU.1 genome wide binding in human primary neutrophils (CD16+ CD66b+) isolated from 93 donors of the BLUEPRINT project. The same donors were previously characterised at genome-wide DNA sequence and multi-level regulatory annotation^7^ (Figure 1a). We identified 36,530 TF-binding peaks across the 93 individuals (Online Methods, Supplementary Figure 1, Supplementary Table 1) and used normalised read counts at peak regions to determine transcription factor quantitative trait loci (tfQTLs; Online Methods). We detected 1,868 independent (linkage disequilibrium [LD] r^2^≥0.8) PU.1 binding QTLs at a False Discovery Rate [FDR] <0.05 (Supplementary Table 2). Lead PU.1 tfQTL SNPs showed a bimodal distribution of distances to their respective differential binding peaks (Figure 1b, Supplementary Table 3), with just over half of them (55%, 1,036/1,868) mapping proximally from the peak edge (<2.5kb; median distance 264bp), and the remaining SNPs (45%; 995/1,868) localising more distally (2.5Kb-1Mb, median distance 23Kb)^21,22^. As shown for other cell types^22^, tfQTL effect sizes were stronger for proximal compared to distal variants (t-test p=2.2×10^−16^, Figure 1c). We further validated a subset of the detected tfQTLs using allele-specific association analysis^23^ (Online Methods), which confirmed a significant allelic imbalance for the majority of the tested peaks (98.8% and 95.5% for peaks associated with proximal and distal variants respectively; Figure 1d).

**Figure 1.**
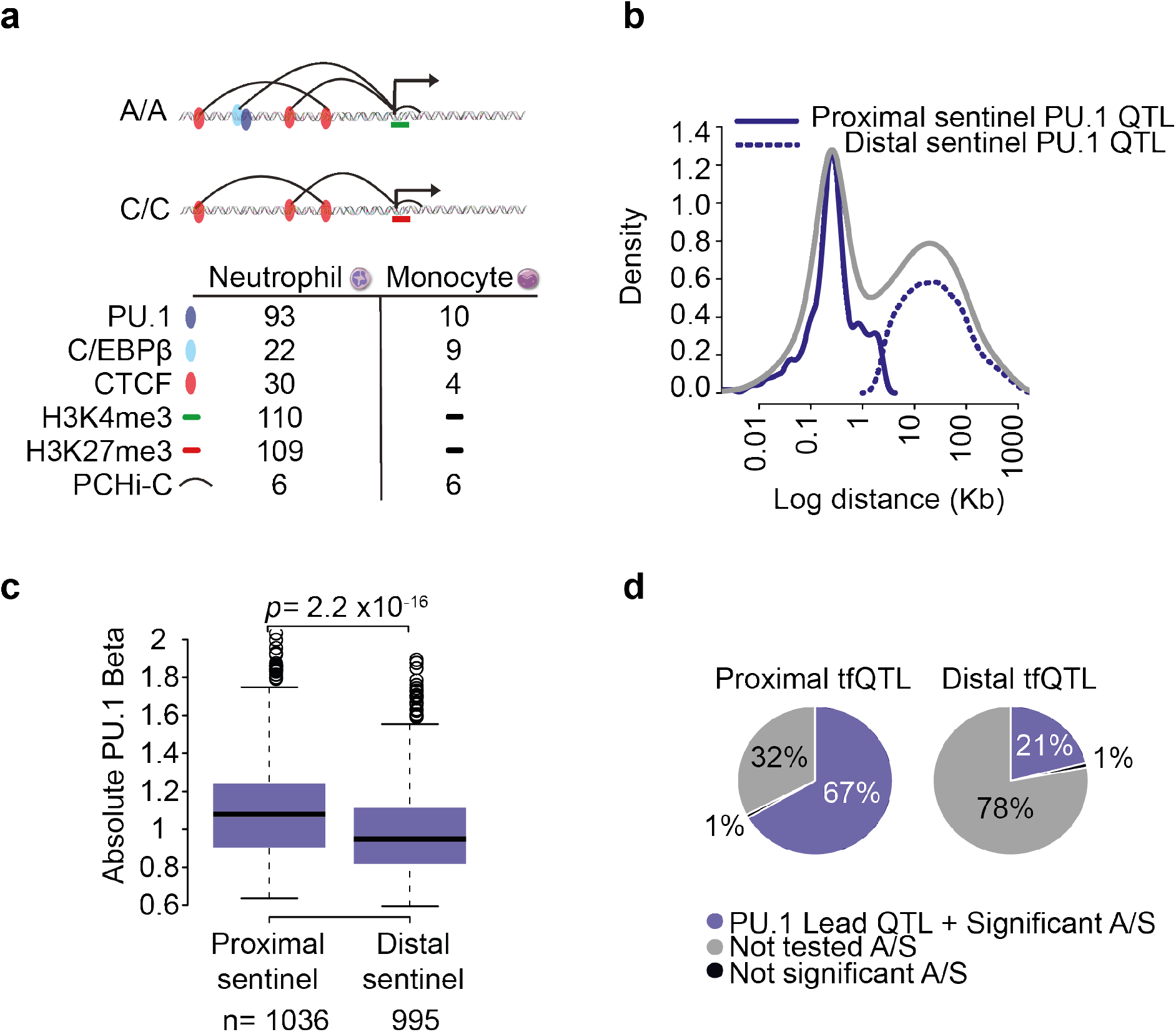
Properties PU.1 transcription factor QTLs. **a**. Summary of molecular traits generated as part of this study. **b**. Density distribution of the distance between sentinel SNPs and their associated PU.1 peaks. The bimodal distribution (grey) can be further subdivided into proximal (solid navy, <2.5Kb) and distal SNP effects (dotted, >2.5Kb). **c**. Boxplot of absolute PU1 tfQTL effect sizes (beta). Proximal PU.1 tfQTLs exhibit larger effect sizes compared to distal tfQTLs (t-test p-value). **d**. Proportion of significant tfQTL SNPs with significant allele-specific (A/S) binding. Peaks without suitable heterozygous SNPs were not tested (grey).

Binding of pioneer TFs to DNA alters local nucleosome positioning, thus allowing recruitment of activating co-factors^24^. However, DNA recognition sequence alone is not sufficient to establish occupancy, and secondary collaborating factors are required to maintain affinity^25^. C/EBPβ is upregulated throughout neutrophil terminal differentiation^18^ and has been shown to co-occupy myeloid enhancers at thousands of PU.1 bound sites^26,27^. The constitutively expressed CTCF is known to play a role in gene regulation by anchoring chromatin interactions^28^, but is not known to functionally associate with PU.1. We assayed these two additional TFs in neutrophils from a subset of overlapping individuals (n=22 donors with QC-pass assays for C/EBPβ, n=30 for CTCF), and identified 18,862 C/EBPβ and 22,197 CTCF filtered peaks from the combined datasets. We performed QTL analysis as before, and prioritised 427 C/EBPβ and 769 CTCF putative tfQTLs reaching a nominal p-value threshold (p≤1×10^−5^; Supplementary Table 2). We found that C/EBPβ tfQTLs effect sizes decreased with increasing distance from PU.1 tfQTLs (Figure 2a-b), reflecting cooperative binding of PU.1 and C/EBPβ at myeloid enhancers^26,29^. Interestingly, CTCF tfQTLs displaying a shared genetic effect with PU.1 predominantly involved CTCF-bound regions located distally to the PU.1 tfQTL lead SNP, suggesting that PU.1 QTL genetic effects may be in part mediated by the 3D chromosomal architecture (Figure 2c).

**Figure 2.**
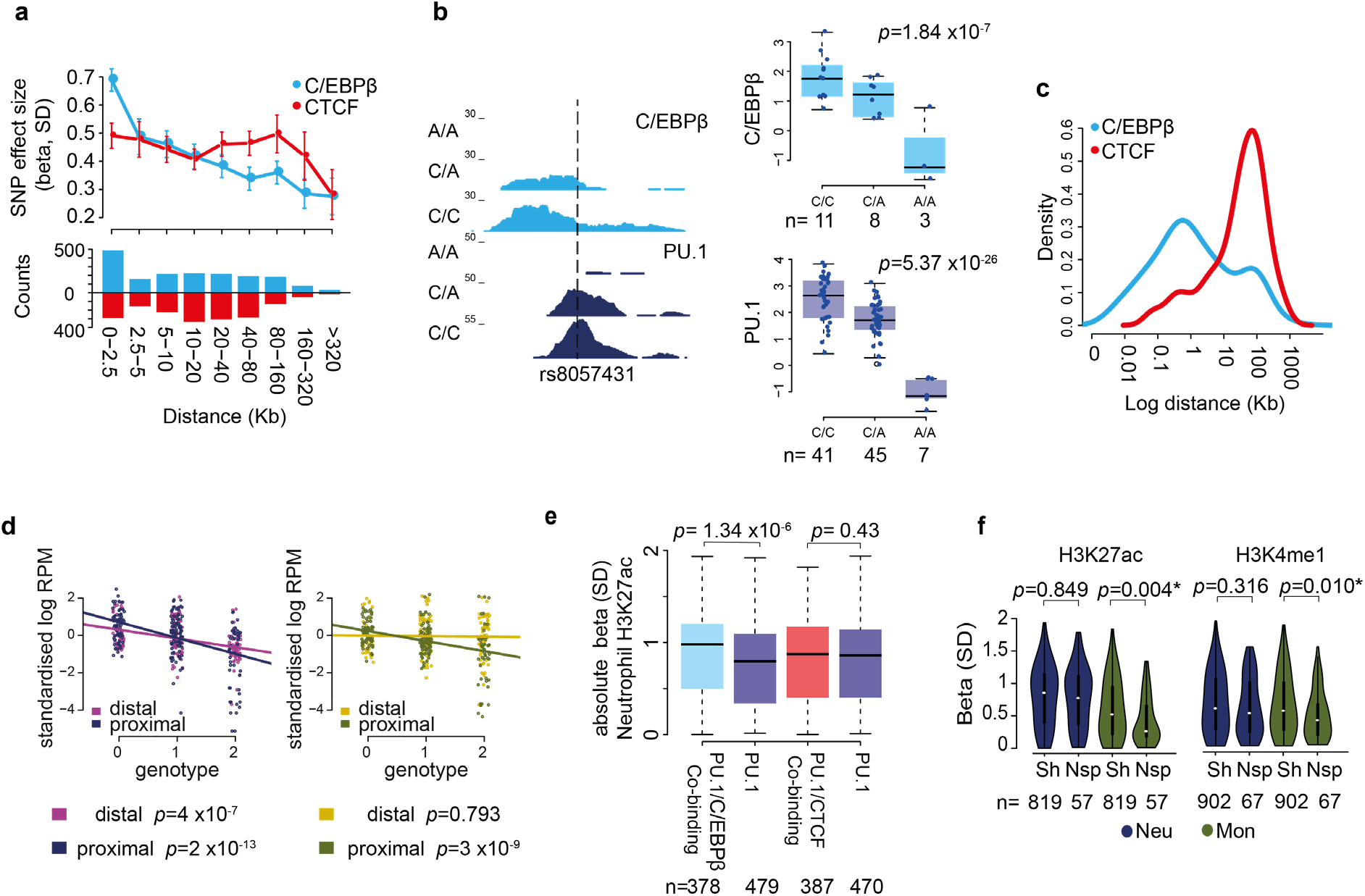
Effect of PU.1 SNPs effect on proximal second transcription factor binding. **a**. Effect size (95% confidence intervals) for association of proximal sentinel PU.1 SNPs with the nearest C/EBPβ (light blue) and CTCF (red) binding site. The effect size decreases with distance for C/EBPβ (linear model p<2.2×10^−16^) but not for CTCF (p = 0.113). Beneath: bar chart of number of peaks included in each distance bin. **b**. Illustrative example of shared tfQTL, where SNP rs8057431 (dashed line) alters a PU.1 motif and is associated with a disruption in binding of both PU.1 and C/EBPβ. Right: signal box plot across all individuals segregated by genotype. Left: raw regional signal of binding intensity for three individuals segregated by genotype. **c**. Density plot of distances of sentinel PU.1 SNP from the nearest C/EBPβ (light blue) or CTCF (red) tested peak (p<10^−5^). **d**. Signal binding intensity (Log RPMs) at PU.1 binding sites normalised by sample at shared PU.1 and C/EBPβ tfQTL sites (33 distal and 43 proximal). Linear models were fitted separately for proximal and distal sites in neutrophils (left panel) and monocytes (right panel) using five matched individuals. **e**. Boxplot of absolute beta for H3K27ac neutrophil QTL (no significance threshold) for proximal tfQTL PU.1, differentiating H3K27ac regions that are or are not marked by C/EBPβ and/or CTCF. **f**. Violin plot showing distribution of effect sizes (Beta) for H3K27ac and H3K4me1 hQTLs for proximal tfQTLs in neutrophils (navy) and monocytes (green), for regions marked by shared (Sh) or neutrophil specific (Nsp) binding. P-values obtained from t-test.

Transcription factor occupancy has been shown to act predominantly through *cis* regulatory SNPs, where coordination of cis-acting variants has been shown to decay with increasing physical distance of SNPs from bound regions^30^. To assess the potential sharing of our tfQTLs across cell types, we additionally generated PU.1 binding maps in primary monocytes (CD14+CD16^−^) isolated from ten BLUEPRINT donors^7^, five of which overlap with the neutrophil PU.1 dataset. Of the neutrophil PU.1 peaks implicating a tfQTL, 93% were also observed in monocytes (Supplementary Figure 2a). The low number of donors tested did not allow us to carry out a tfQTL analysis in monocytes. To assess coordination of genetic effects at PU.1 binding sites across cell types, we therefore assessed the strength of binding at monocyte peaks for individuals stratified by PU.1 tfQTL lead variant genotype. We found the monocytes displayed consistent direction and strength of binding at proximal SNPs (linear regression *p*=3×10^−9^) compared to neutrophils (*p*=2×10^−13^), compatible with shared genetic effects between the two cell types. However, the same was not true for distal SNPs (neutrophils p=4×10^−7^, monocytes p=0.793; Figure 2d), which may be driven by more complex and cell-type specific long-distance chromatin contacts.

To explore coordination of genetic influences on PU.1 binding and local chromatin state, we initially took advantage of the previously published histone associated QTL (hQTL) data for the enhancer-associated histone marks H3K4me1 and H3K27ac in neutrophils^7^. In total, 808 H3K4me1 and 946 H3K27ac lead hQTL SNPs overlapped (r^2^≥0.8) PU.1 tfQTLs. We next generated binding profiles of the active promoter-associated histone mark H3K4me3 and Polycomb-associated repressive mark H3K27me3 in neutrophils (n=110 and n=109 donors, respectively) identifying 621 and 367 shared tfQTL/hQTLs, respectively (Supplementary Table 2). Using the pi1 statistic^31^, we found evidence of sharing between PU.1 tfQTLs with hQTLs in both neutrophils and monocytes (pi1_H3K27ac_=0.73-0.76, and pi1H3K4me1=0.76-0.80). Sharing between neutrophil PU.1 tfQTLs and hQTLs detected in CD4 naïve T cells was lower (pi1_H3K27ac_=0.36-0.72, and pi1_H3K4me1_=0.30-0.79; Supplementary Figure 2c), compatible with PU.1 not being expressed in the latter (Supplementary Figure 2c)^32^. Further, H3K27ac marked regions^7^ co-occupied by PU.1 and C/EBPβ displayed greater hQTL effect sizes compared to peaks bound by PU.1 alone (t-test *p*=1.34×10^−6^; Figure 2e), suggesting stronger genetic effects for enhancers at co-occupied sites in neutrophils^29^. Consistent with this, cell type-specific binding of PU.1 and C/EBPβ correlated with cell type-specific chromatin activity (Supplementary Figure 3a-b). H3K27ac and H3K4me1 hQTLs intersecting proximal neutrophil-specific PU.1 tfQTLs had significantly lower effect sizes in monocytes compared to cell-shared sites (Figure 2f), consistent with a neutrophil-specific role of PU.1 in activating chromatin state in these regions.

We next assessed the distance between the PU.1 and histone mark peaks for each shared tfQTL-hQTL genetic association. As previously observed^21^, there was a pronounced bimodal distribution of distances between PU.1 binding peaks and the locations of H3K27ac and H3K4me3 marks (Figure 3a), with around a half of PU.1 peaks localizing to less than 1kb away from the respective H3K27ac and H3K4me3 peaks, and others mapping 10-100kb from them. Given that H3K4me3 is associated with active promoters, this observation highlights the potential long-range regulatory effects of PU.1 binding to distal DNA elements on promoter activity, which are commonly mediated by three-dimensional DNA looping interactions^33^. To investigate the role of PU.1 long distance regulation, we generated Capture Hi-C (PCHi-C) profiling in neutrophils and monocytes isolated from three donors each and integrated these data with previously published PCHi-C data for these cell types in three more individuals^34^ (Supplementary Figure 4a-b). We detected ~190,000 Promoter Interacting Regions (PIRs) in total across neutrophils and monocytes (CHiCAGO score > 5)^35^, ~82,000 of which were detectable in each of the cell types (Supplementary Figure 4c). PIRs enriched in PU.1 binding and enhancer-associated H3K4me1/H3K27ac marks were correlated with the level of expression of the genes they contacted (Figure 3b), as previously shown in the context of other cell types^36–38^. In contrast, CTCF binding at PIRs did not correlate with target gene expression (Figure 3b), as expected given the constitutive nature of many CTCF-mediated chromosomal interactions. Notably, the PIRs of genes showing differential expression between neutrophils and monocytes were enriched (100 permutations, *p*≤0.01) for the binding of PU.1 and C/EBPβ in the highest-expressing cell type (Figure 3c). Consistently we also found cell type-specific binding of these TFs to be enriched within cell type-specific PIRs (permutation *p*≤0.01) (Figure 3d). Jointly, these results reinforce the role of PU.1 and C/EBPβ in establishing tissue-specific transcriptional patterns.

**Figure 3.**
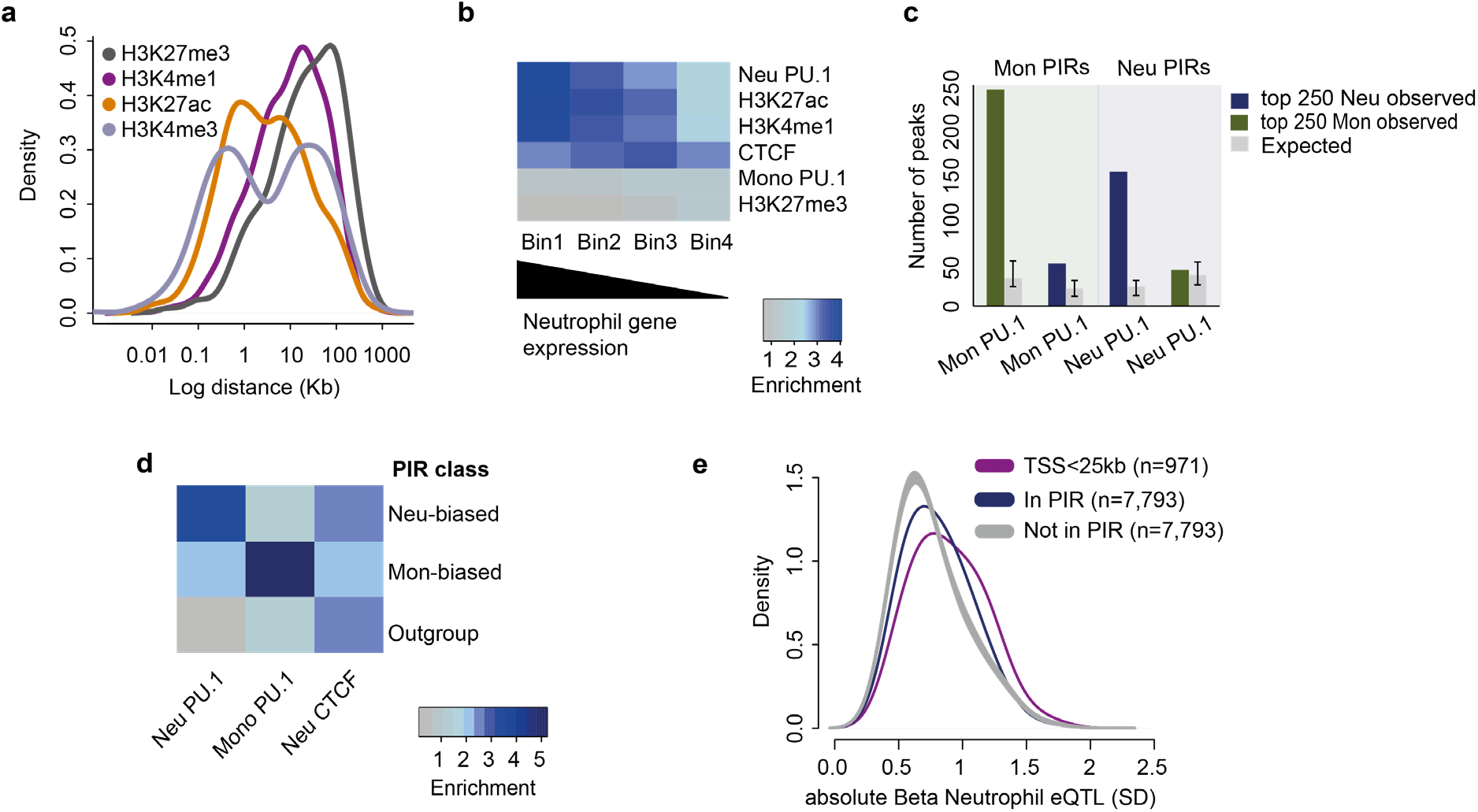
PU.1 tfQTLs mediate gene regulation through chromatin contacts. **a**. Density distribution plot of log distance between lead PU.1 tfQTL SNPs and shared (r_2_≥0.8) lead histone QTL peaks in neutrophils. **b**. Heat map showing enrichment of transcription factor or histone modification regions intersecting PIRs, whereby PIRs were ranked into four bins based on the gene expression of connected baited genes in neutrophils. **c**. Bar plot of the number of cell type specific TF binding sites intersecting PIRs in four scenarios derived from PCHi-C datasets from monocytes and neutrophils (left and right hand panels respectively), and for PIRs connected to the top 250 differentially upregulated genes in monocytes (green bars) or in neutrophils (blue bars; see also Supplementary Figure 3). Grey bars indicate the number of TF intersecting PIRs for randomly shuffled transcription factor binding sites. **d**. Heat map showing enrichment of DiffBind PU.1 regions in neutrophils and monocytes (Supplementary Figure 3), and neutrophil CTCF. Overlapping PIRs were classified into three categories of cell type specificity (see Methods). **e**. Density plot of gene expression QTL Beta for neutrophil (navy) PU.1 SNPs within PIRs versus distance-matched significant SNPs not in PIRs (grey) (t-test p<2×10^−16^), and distribution of betas for SNPs within <25Kb of transcription start site (purple). The SNPs that are not in PIRs are also significant PU.1 tfQTLs and eQTLs (p<1×10^−5^ cut-off).

We next investigated the effect of PU.1 binding variation at PIRs on the expression of target genes. PU.1 tfQTLs were intersected with PCHi-C and expression QTL (eQTL) data from the Chen *et al.* study^7^. Only PU.1 tfQTL/eQTL (p<1×10^−5^) pairs located distally to TSSs (>25Kb) were considered, in order to exclude eQTLs implicating promoter-based variants and ensure a high resolution of PCHi-C signal detection. PU.1 tfQTL SNPs mapping to PIRs showed significantly larger effects on the expression of the genes they contacted compared with distance-matched SNPs that did not map to a PIR (t-test p<2×10^−16^; Figure 3e), in agreement with physical interactions playing a role in mediating the distal regulatory effects of PU.1 binding.

To explore the extent to which genetic variation affecting PU.1 binding may directly affect promoter-enhancer interactions, we employed an allele-specific strategy (Methods) to identify heterozygous sites within PIRs that exhibited allelic imbalance at PCHiC contacts (Supplementary Figure 4d-e). We found that ~14,000 heterozygous SNPs within PIRs that displayed evidence of allelic bias in both neutrophils and monocytes were enriched for PU.1 and CTCF binding (Figure 4a-c, Supplementary Figure 4f). Notably, the same was true for the hQTLs for the Polycomb-associated inhibitory mark H3K27me3, consisted with a role of Polycomb repressive complexes in shaping regulatory chromatin architecture^39^. An example of a SNP showing allelic imbalance affecting promoter-enhancer connectivity was rs519989, which was also associated with PU.1 binding, histone modifications and expression the gene *LRRC8C* (Figure 4d-f; Supplementary Table 4). *LRRC8C* encodes a volume-regulated anion channel subunit^40^ upregulated during terminal differentiation of neutrophils^41^. This and other loci thus demonstrate coordinated genetic influences on PU.1 binding, chromatin activity and the formation of promoter interactions in the regulation of neutrophil gene expression.

**Figure 4.**
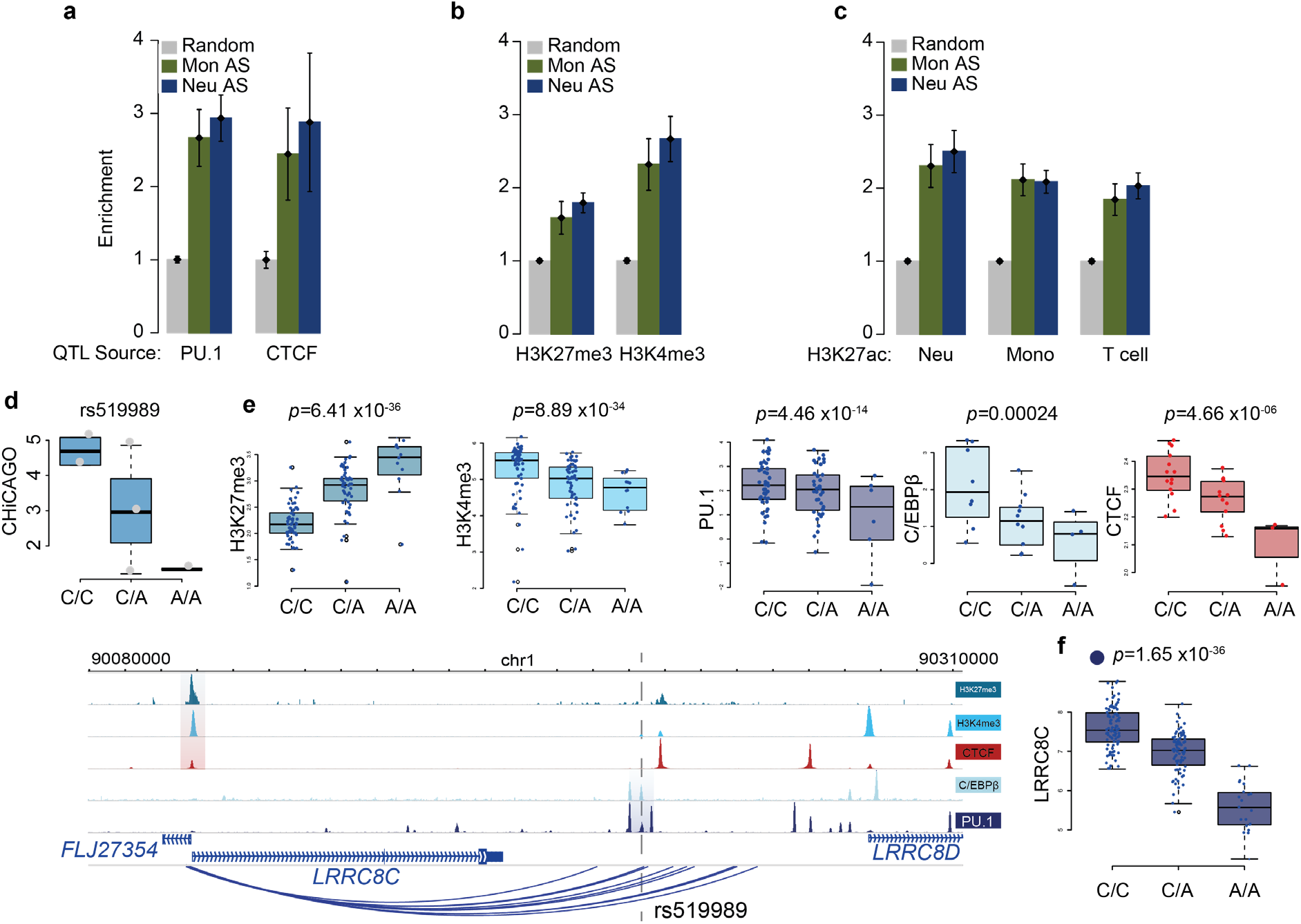
tfQTLs perturb gene expression through altered chromatin state. **a**. Enrichment of significant tfQTLs (PU.1 and CTCF; p<1e-5), **b**. significant hQTLS (H3K27me3, H3K4me3; p<1e-5), **c**. and significant hQTLs (H3K27ac; p<1e-5) in PIRs. K27AC-Mon represents significant QTLs found in monocytes (p<1e-5) and tested against PIRs found in both monocytes and neutrophils. Similarly, H3K27ac-Neu and H3K27ac-Tcell represent significant QTLs (p<1e-5) found in neutrophils and T-cells and tested against PIRs found in both monocytes and neutrophils. **d**. CHiCAGO scores for PIR at lead QTL rs519989 segregated by donor genotype. **e**. Top; Signal boxplots with donors separated by genotype for rs519989 for five molecular traits. Below; Genome browser view of region around *LRRC8C* gene, QTL regions for each molecular trait are highlighted. Dashed line depicts location of rs519989. **f**. Boxplots of RNA level for *LRRC8C* gene segregated by donor genotype for SNP rs519989.

Finally, to explore the influence of the identified PU.1 tfQTLs and their potential downstream effects on haematological traits and diseases, we accessed summary statistics from public GWAS studies of cell-matched full blood count traits^2^ and autoimmune diseases^42–47^ PU.1 binding regions were enriched^48^ for GWAS SNPs associated with myeloid cell traits (eg. neutrophil counts) and with autoimmune diseases (Figure 5a). We formally tested the overlap of PU.1 tfQTLs and GWAS SNPs using colocalisation analysis^49,50^, revealing 43 proximal and 74 distal tfQTLs that shared a genetic signal (posterior probability [PP]>0.9) with at least one GWAS locus (Table 1, Supplementary Table 5). We next used CATO (Contextual Analysis of TF Occupancy)^51^ to identify PU.1 collaborating factors that may be involved in mediating these traits at shared PU.1 tfQTL / GWAS loci. Colocalising SNPs were shown to affect predominantly binding recognition motifs for several PU.1 binding partners, including C/EBP, AP-1, ETS, CTF/NF-1, ATF/CREB and RUNX (Supplementary Figure 5)^52^. These results highlight the likely role of PU.1 and its partners in mediating the functional effects of GWAS variants in neutrophils.

**Figure 5.**
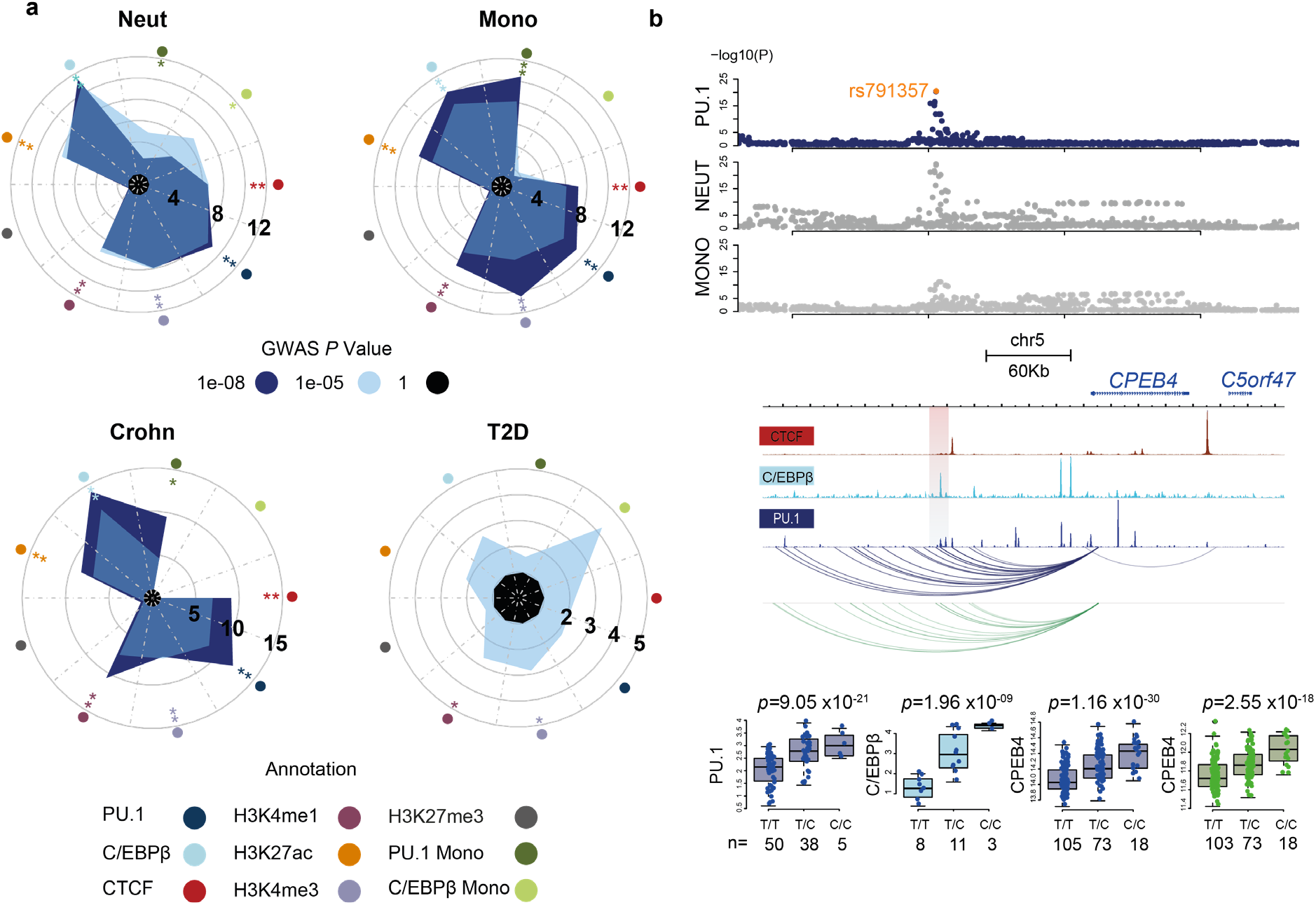
tfQTLs influence cellular phenotype and disease. **a**. Circos plots displaying fold-enrichment of GWAS loci within functional annotations derived fromTF binding and histone modifications in neutrophils and DiffBind derived peaks for PU.1 and C/EBPβ in monocytes. Four illustrative GWAS summary statistics were used: neutrophil count, monocyte count, Crohn’s disease and Type 2 diabetes (T2D) as negative control. Radial grid lines for GWAS p-values, asterisk denotes significance of enrichment for annotation tested at each GWAS p-value cut off. **b**. Example of colocalised signal for sentinel SNP rs791357 which is significant GWAS with a shared association for both neutrophil and monocyte count traits. Top; Manhattan plots showing p-value distribution for shared SNPs neutrophil and monocyte counts and PU.1 tfQTL (navy). Middle; genome visualisation of TF binding for CTCF, C/EBPβ and PU.1. The top associated peak is highlighted by the shaded area. *CPEB4* baited PIRs for both neutrophils (blue) and monocytes (green). Bottom; boxplot for TF and RNA signal segregated by donor genotype. PU.1 (navy), CEBPβ (light blue), CPEB4 gene expression neutrophil (navy) and CPEB4 gene expression monocyte (olive green). rs791357 associated with PU.1 in neutrophils. In addition, PCHiC data shows that this region is highly connected to the enhancer region in both neutrophils and monocytes.

**Table 1.**
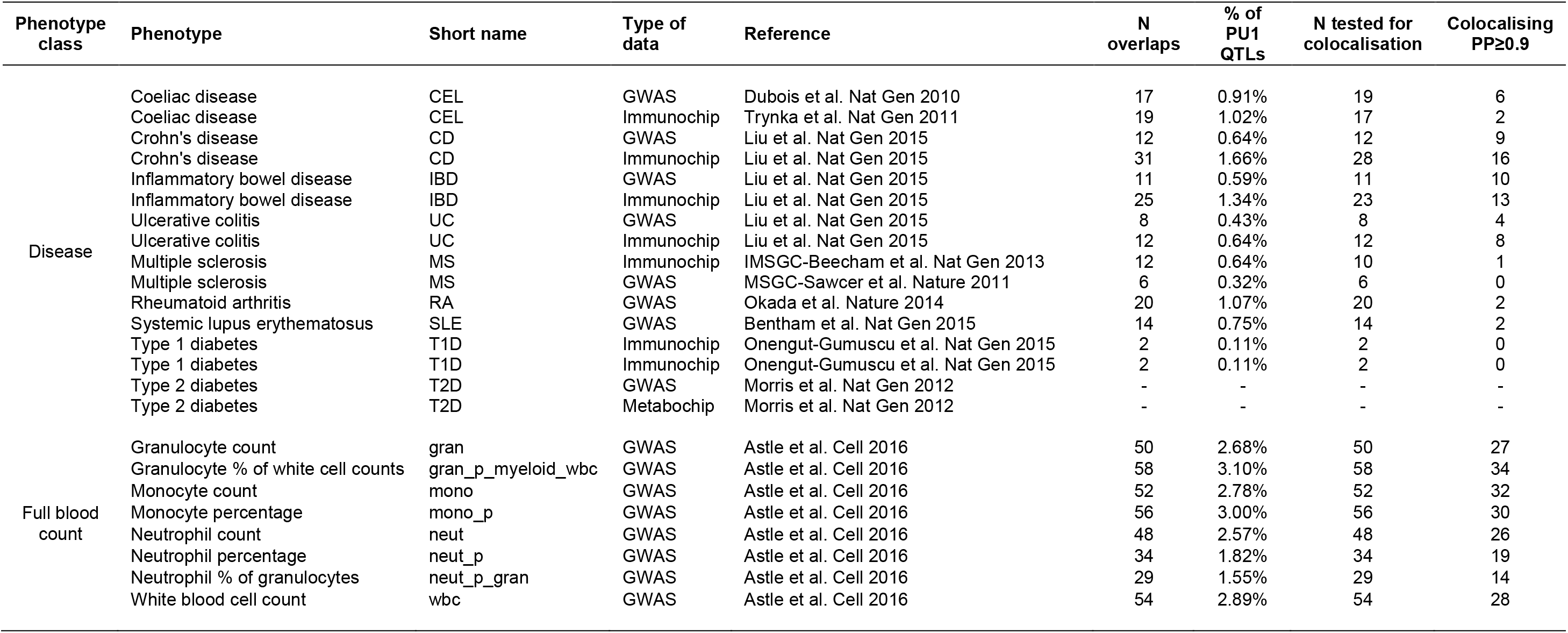
Overview of PU1 tfQTL overlap with disease and blood cell count loci. Summary table of the numbers of colocalised loci for PU.1 tfQTL and tested GWAS summary statistics.

To determine the putative target genes underpinning PU.1-mediated disease associations, we integrated PCHi-C and eQTL data^7^ in neutrophils. Overall, 27 high confidence target genes at QTL loci colocalised with GWAS summary statistics (Supplementary Table 6). Interestingly, 35% of the shared tfQTL / GWAS SNPs could be attributed to a proximal tfQTL at a PU.1 binding site. This finding suggests that many of PU.1 tfQTL themselves are under distal genetic control potentially mediated through enhancer-enhancer interactions^53–55^. One such example is the rs791357_C variant associated with decreased neutrophil and monocyte cell counts. PCHi-C data shows that this region is highly connected to the *CPEB4* gene in both neutrophils and monocytes (Figure 5b). CPEB4 is a cytoplasmic polyadenylation element which binds to recognition sequence in PolyA tail of mRNAs and can activate or inhibit translation^56^. CPEB4 is involved in controlling terminal differentiation in erythroid cells^57^ and the proliferation of some cancers^58^. The SNP rs791357 is a proximal tfQTL for PU.1 (*p*=9.05×10^−21^) and C/EBPβ (*p*=1.963×10^−9^), an hQTL for H3K4me3 (*p*=1.98×10^−17^) and H3K27ac (*p*=1.41×10^−33^) and an eQTL for *CPEB4* (*p*=1.16×10^−30^) in neutrophils. Similar sharing of PU.1 and C/EBPβ binding site was observed in monocytes with hQTL (H3K27ac *p*=8.14×10^−26^) and eQTL for *CPEB4* (*p*=2.55×10^−18^). Additional example loci shared tfQTL function through multiple traits and the presence of PCHi-C interactions between enhancers and colocalised genes (Supplementary Figure 6a-b).

In conclusion, our analysis suggests that genetically-determined variation in PU.1 binding in neutrophils modulates gene expression, acting via changes in the local chromatin state and, at least in some cases, in the patterns of promoter-enhancer interactions. We show that these effects underpin the genetic associations for a number of important human blood cell traits and diseases, confirming the role of PU.1 in neutrophil biology and implicating this cell type as a potentially causal for a number of autoimmune traits.

## Author Contributions

Conceived and designed the study, S.W., B.M.J., M.S., and N.S. Performed experiments, S.W., and B.M.J. Generated experimental resources, F.B., S.F., and B.F. Performed formal analysis, S.W., L.V., A.L.M., K.K., L.C., Y.Y., S.E., V.I., H.E., M.T., D.R., A.D., and M.S. Investigation, S.W., L.V., K.W., and A.L.M. Data Curation, L.V., Y.Y., H.P., D.R., A.D., and L.C. Supervision and study coordination, P.F., L.C., K.D., T.P., P.F., M.F., M.S., and N.S. Project Administration, D.M., L.C., K.D., P.F., M.F., M.S., and N.S. Performed primary manuscript writing S.W., A.L.M., M.S., and N.S.

## Competing Interests statement

The authors declare no competing interests.

## Online Methods

### Sample collection and cell isolation

#### Peripheral adult blood collection

ChIP-seq data generated in this study used donor samples which were collected as part of the previously described study^7^. Blood was obtained from donors who were members of the NIHR Cambridge BioResource (http://www.cambridgebioresource.org.uk/) with informed consent (REC 12/EE/0040) at the NHS Blood and Transplant, Cambridge. Donors were on average 55 years old (range 20-75 years old), with 46% of donors being male. A unit of whole blood (475 ml) was collected in 3.2% Sodium Citrate. An aliquot of this sample was collected in EDTA for genomic DNA purification. A full blood count (FBC) for all donors was obtained from an EDTA blood sample, collected in parallel with the whole blood unit, using a Sysmex Haematological analyser. The level of C-reactive protein (CRP), an inflammatory marker, was also measured in the sera of all individuals. All donors used for the collection had FBC and CRP parameters within the normal healthy range. Blood was processed within 4 hours of collection.

#### Isolation of cell subsets

Samples were as those as described in^7^. To obtain pure samples of ‘classical’ monocytes (CD14+ CD16-) and neutrophils (CD66b+ CD16+) we implemented a multi-step purification strategy. Whole blood was diluted 1:1 in a buffer of Dulbecco’s Phosphate Buffered Saline (PBS, Sigma) containing 13mM sodium citrate tribasic dehydrate (Sigma) and 0.2% human serum albumin (HSA, PAA) and separated using an isotonic Percoll gradient of 1.078 g/ml (Fisher Scientific). Peripheral blood mononuclear cells (PBMCs) were collected and washed twice with buffer, diluted to 25 million cells/ml and separated into two layers, a monocyte rich layer and a lymphocyte rich layer, using a Percoll gradient of 1.066g/ml. Cells from each layer were washed in PBS (13mM sodium citrate and 0.2% HSA) and subsets purified using an antibody/magnetic bead strategy. To purify monocytes, CD16+ cells were depleted from the monocyte rich layer using CD16 microbeads (Miltenyi) according to the manufacturer’s instructions. Cells were washed in PBS (13mM sodium citrate and 0.2% HSA) and CD14+ cells were positively selected using CD14 microbeads (Miltenyi). To purify neutrophils, the dense layer of cells from the 1.078 g/ml Percoll separation was lysed twice using an ammonium chloride buffer to remove erythrocytes. The resulting cells (including neutrophils and eosinophils) were washed and neutrophils positively selected using CD16 microbeads (Miltenyi) according to the manufacturer’s instructions. The purity of each cell preparation was assessed by multicolour FACS using conjugated antibodies for CD14 (MφP9, BD Biosciences) and CD16 (B73.1 / leu11c, BD Biosciences) for monocytes, CD16 (VEP13, MACS, Miltenyi) and CD66b (BIRMA 17C, IBGRL-NHS) for neutrophils. Purity was on average 95% for monocytes and 98% for neutrophils.

#### ChlP-sequencing

Purified cells were fixed with 1% formaldehyde (Sigma) at a concentration of approximately 10 million cells/ml. Fixed cell preparations were washed and stored re-suspended in PBS at 4°C prior to lysis and sonication. Sonication protocols were performed in a Diagenode PicoRuptor for 8 cycles of 30 seconds on, 30 seconds off in a 4°C water cooler. Samples were checked for sonication efficiency using the criteria of 150-500bp, by Agilent DNA bioanalyzer. ChIP-seq was carried out as previously described^59^ all liquid handling steps were performed on an Agilent Bravo NGS. Protein A Dynabeads (Invitrogen) were coupled with 2.5μg of antibody. Sonicated lysate (3-5 million cells) was then added to the bead/antibody mix and incubated at 4°C overnight. ChIP-DNA bound beads were washed for ten repetitions in cold RIPA solution. Elution of DNA from beads at 65°C for five hours to reverse the cross linking process. 2μl RNase was added to ChIP-DNA and incubated at 37°C for 30 minutes, followed by 2μl of Proteinase K treatment at 55 °C for 1 hour. 1:1.8 ratio of Ampure beads (Beckman Coulter, A63881) were added to the DNA followed by two cold 70% ethanol washes. ChIP-DNA was eluted in 50μl elution buffer. Illumina sequencing libraries were prepared on a Beckman Fx liquid handling system. End-repair, A-tailing and paired-end adapter ligation were performed using NEBnext reagents from New England Biolabs (E6000S), with purification using a 1:1 ratio of AMPure XP to sample between each reaction. Amplification of ChIP-DNA was performed using Kapa HiFi master mix (Kapa Biosystems KK2602), 18 cycles of PCR followed by a 0.7:1 Ampure XP clean-up. Antibodies for H3K4me3 (C15410003), H3K27me3 (C15410195), CTCF (C15410210) were obtained from Diagenode, Liege, Belgium. Antibodies for PU.1 (sc-352x, sc-22805x) and C/EBPβ (sc-150x) were obtained from Santa Cruz Biotechnology.

#### Data processing and peak calling

ChIP libraries were sequenced using Illumina HiSeq 2000 and HiSeq 2500 at 50bp single end reads. Sequenced reads were aligned to reference genome using BWA (bwa *aln −q* 15). Duplicate reads were marked using Picard MarkDuplicates (v1.103). Reads with mapping quality less than 15 were removed (SAMtools v0.1.18). The fragment size L for each aligned bam was estimated using PhantomPeakQualTools vr18, which uses cross correlation of binned read counts between forward and reverse strands. To identify highly enriched genomic regions, we used MACS2^60^ (v2.0.10.20131216, standard options) for peak calling with the estimated fragment size from PhantomPeakQualTools *(--shiftsize=half fragment size),* with narrow for PU.1, C/EBPβ, CTCF, H3K4me3 and broad flags set for H3K27me3. For background control ChIP input was created from merging random selected samples. Reads from 4 pools of 12 individuals for neutrophil input and 2 pools of 6 individuals for monocytes. ChIP inputs were as follows:

| ID | Samples | Cell Type | Gender |
| --- | --- | --- | --- |
| NS1140 | pool of S00W29, S00WP0, S00FK4 | Neutrophil | Female |
| NS1141 | pool of S00JT7, S00HVB, S00M0G | Neutrophil | Male |
| NS1163 | pool of S00T4, S00NXK, S00PBJ | Neutrophil | Female |
| NS1164 | pool of S00RMQ, S00RD7, S00NRW | Neutrophil | Male |
| NS1556 | pool of S00W29, S00WP0, S00KHR | Monocyte | Female |
| NS1557 | pool of S00GBI, S00JV3, S00M2C | Monocyte | Male |

Significant peaks were selected to be at 1% FDR or less.

#### Data Quality

We removed ChIP samples that had a relative strand correlation (RSC) < 0.8 and normalised strand correlation (NSC) < 1.05^61^. We defined high confidence data those from ChIP with RSC > 0.8 and NSC > 1.05. Otherwise, we used genome browser tracks to confirm visually a good ChIP and include it in the final data set. Supplementary Figure 1 and Supplementary Table 1 shows quality control metrics and corresponding principal components, showing no batch effects after PEER correction using K=10 factors.

#### Normalised read count in the reference peak set

Consensus peak sets were constructed using dba.peakset function within DiffBind R package^62,63^. http://bioconductor.org/packages/release/bioc/vignettes/DiffBind/inst/doc/DiffBind.pdf. For PU.1, H3K4me3 and H3K27me3 we set the minimum number of samples for a peak to be included in consensus to 3, for C/EBPβ, CTCF and monocyte samples minimum was set to 2. Sex chromosomes were not included in the QTL analysis. The reference peak set was filtered further for read counts as described below. Next, we generated quantification signal of ChIP-seq for each donor. Here we only considered read counts under the peaks, as the regions outside peaks are more likely to be noise or background signal than true enrichment. For each donor, we generated a vector of log2 reads per million (log2RPM) per peak in the reference peak set by counting the number of overlapping reads under the peaks (BEDOPS bedmap-count) and normalised the counts with the total number of reads in the library. We further filtered the reference peak set to only consider peaks with log2RPM > 0 in at least 50% of the donors in a given cell type, corrected for ten PEER factors and applied quantile normalisation across donors. For QTL calling with H3K27me3, two sets of summary statistics are provided on two separate signal matrices. In the first set H3K4me3 peak annotations were used in conjunction with H3K27me3 signal to enrich for poised promoter QTLs. In the second set broad called H3K27me3 peaks were divided into 2500bp windows.

#### Identification of PU.1 and C/EBPβ differential binding sites

We used DiffBind version 1.12.0 with default EdgeR (3.8.3) option to identify peaks which were differentially bound between neutrophils and monocytes. We used the six best quality samples and their peak sets for this analysis:

| ID | Individual | Factor | Cell Type |  | ID | Individual | Factor | Cell Type |
| --- | --- | --- | --- | --- | --- | --- | --- | --- |
| NS1509 | S00QTG | PU.1 | Monocyte | | NS1559 | S00V57 | CEBPβ | Monocyte |
| NS1510 | S00M48 | PU.1 | Monocyte | | NS1565 | S0100M | CEBPβ | Monocyte |
| NS1511 | S00PDF | PU.1 | Monocyte | | NS1563 | S00U3F | CEBPβ | Monocyte |
| NS1514 | S00HVB | PU.1 | Monocyte | | NS1558 | S00TT4 | CEBPβ | Monocyte |
| NS1516 | S00GBI | PU.1 | Monocyte | | NS1562 | S00XHC | CEBPβ | Monocyte |
| NS1522 | S00YEE | PU.1 | Monocyte | | NS1566 | S00U1J | CEBPβ | Monocyte |
| NS1463 | S011HL | PU.1 | Neutrophil | | NS585 | S00DP2 | CEBPβ | Neutrophil |
| NS1464 | S013HD | PU.1 | Neutrophil | | NS743 | S00NHF | CEBPβ | Neutrophil |
| NS1554 | S00WKA | PU.1 | Neutrophil | | NS791 | S00QQM | CEBPβ | Neutrophil |
| NS1551 | S00SMM | PU.1 | Neutrophil | | NS729 | S00M2C | CEBPβ | Neutrophil |
| NS1437 | S00YK2 | PU.1 | Neutrophil | | NS717 | S00K4G | CEBPβ | Neutrophil |
| NS1490 | S00YEE | PU.1 | Neutrophil | | NS793 | S00QWA | CEBPβ | Neutrophil |

We selected peaks present in at least three individuals and that had a minimum three-fold difference in binding signal as cut off. Heatmap visualisation of differentially bound regions Deeptools 2^64^.

#### Transcription factor enrichments

For determining enrichment of ChIP-seq regions of interest within PIRs we used regioneR (1.0.3)^65^, which performs a statistical evaluation of two sets of genomic regions by permutation testing. We set to 50 permutations the randomisation of genomic regions to determine the null. In Figure 3b-d are Neu PU.1 and Mono PU.1 regions identified from DiffBind differential binding analysis (Supplementary Figure 3a). In Figure 3D are cell type biased PIRs were constructed using data from Javierre et al. We took a subset PIRs from B cells, CD8 T cells, CD4 T cells, Neutrophils, Monocytes, Megakaryocytes and Erythrocytes. These were split into three classifications: (i) PIRs that were found in neutrophil and one other cell type, (ii) PIRs that were found in monocyte and one other cell type, and (iii) as an outgroup, megakaryocyte PIRs that were not shared with neutrophil and monocyte.

#### Differentially expressed genes and gene expression counts

Gene expression counts and list of differentially expressed genes were available from Ecker et al.^3^

#### QTL mapping

Cis-acting QTL mapping was done using the LIMIX package^66^, available from github (https://github.com/PMBio/limix). We considered genetic variants mapping to within 1 Mb (on each side) of each tested feature (peak), and tested their association using linear regression. Models were fit on quantile-normalized PEER residuals, also including a random effect term accounting for polygenic signal and sample relatedness (as in the variance component models above we used the realized relatedness matrix to capture sample relatedness). From the linear regression we obtained the effect size and p-value for each tested association. To correct for multiple hypothesis testing, we performed a two-step procedure^67^: first, we corrected for multiple testing across variants for each molecular outcome using Bonferroni correction and, second, we adjusted the obtained p-values for multiple-testing across phenotypes within each layer using a the Q-value procedure^31^, considered QTLs at a significance threshold of 5% FDR.

#### Promoter Capture HiC (PCHi-C)

Cells were isolated as described^34^. One donor was used for preparing each PCHi-C library. In total, twelve PCHi-C libraries were prepared, six using monocytes and six using neutrophils. Approximately 8×10^−7^cells per library were resuspended in 30.625 ml of DMEM supplemented with 10% FBS, and 4.375 ml of formaldehyde was added (16% stock solution; 2% final concentration). The fixation reaction continued for 10 min at room temperature with mixing and was then quenched by the addition of 5 ml of 1 M glycine (125 mM final concentration). Cells were incubated at room temperature for 5 min and then on ice for 15 min. Cells were pelleted by centrifugation at 400g for 10 min at 4°C, and the supernatant was discarded. The pellet was washed briefly in cold PBS, and samples were centrifuged again to pellet the cells. The supernatant was removed, and the cell pellets were flash frozen in liquid nitrogen and stored at −80 °C. Biotinylated 120-mer RNA baits were designed to the ends of HindIII restriction fragments overlapping Ensembl-annotated promoters of protein-coding, noncoding, antisense, snRNA, miRNA and snoRNA transcripts^37^. A target sequence was accepted if its GC content ranged between 25% and 65%, the sequence contained no more than two consecutive Ns and was within 330 bp of the HindIII restriction fragment terminus. A total of 22,076 HindIII fragments were captured, containing a total of 31,253 annotated promoters for 18,202 protein coding and 10,929 non-protein genes according to Ensembl v75 (http://grch37.ensembl.org). Hi-C library generation was carried with in-nucleus ligation as described previously^68^. Chromatin was then de-crosslinked and purified by phenol:chloroform extraction. DNA concentration was measured using Quant-iT PicoGreen (Life Technologies), and 40 μg of DNA was sheared to an average size of 400 bp, using the manufacturer’s instructions (Covaris). The sheared DNA was end-repaired, adenine-tailed and double size-selected using AMPure XP beads to isolate DNA ranging from 250 to 550 bp. Ligation fragments marked by biotin were immobilized using MyOne Streptavidin C1 DynaBeads (Invitrogen) and ligated to paired-end adaptors (Illumina). The immobilized Hi-C libraries were amplified using PE PCR 1.0 and PE PCR 2.0 primers (Illumina) with 7 PCR amplification cycles. PCHi-C. Capture Hi-C of promoters was carried out with SureSelect target enrichment, using the custom designed biotinylated RNA bait library and custom paired-end blockers according to the manufacturer’s instructions (Agilent Technologies). After library enrichment, a post capture PCR amplification step was carried out using PE PCR 1.0 and PE PCR 2.0 primers with 4 PCR amplification cycles. For more details, see^36^. PCHi-C libraries were sequenced on the Illumina HiSeq2500 platform. 3 sequencing lanes per PCHi-C library.

HICUP and CHiCAGO Sequencing reads were processed and mapped with HiCUP and PCHi-C interaction was called using CHiCAGO with default parameters^35,69^.

#### Datasets

Data generated in this study was deposited to the European Genome-phenome Archive under the following accession IDs: transcription factor data: EGAD00001004571; H3K4me3: EGAD00001002711; H3K27me3: EGAD00001002712; PCHiC: EGAS00001001911.

#### Genotyping check of ChlP-Seq and PCHi-C bams

Identity matching for each sample and for each analysis was performed by extracting genotypes from RNA-seq and ChIP-seq and comparing them to SNPs from the WGS data. The first stage of verifying the sample identity concordance between the RNA-seq/ChIP-seq and WGS data involved pre-processing the BAM files for one autosomal chromosome (chr1) to remove PCR duplicates and reads with mapping quality score <10. The variants were then called from the resulting BAM file using *mpileup* from the SAMtools package^70^. The variants with QUAL <20, DP <5 and GQ <5 were filtered out. Then, we compared genotypes of the filtered variants with genotypes generated from WGS and imputation. The genotypes generated were considered to be from the same sample if the concordance rate was greater than 90%.

#### Allele specific analysis of transcription factor binding

For allele specific analysis, we used the phased WGS VCF that was also utilised for QTL mapping but here we removed indels and only considered biallelic single nucleotide variants. We then mapped deduplicated ChIP-seq reads on each allele of each SNVs using GATK ASEReadCounter with default parameters, base quality ≥2 and mapping quality ≥15. We then filtered for heterozygous SNVs only with ≥10 read counts per site and nonzero counts in both alleles. We required 2 donors meeting these read counts criteria at each site. To carry out association analysis, we used Rasqual^23^ with total read counts per sample as offset parameter. Note that Rasqual uses a model that corrects for reference mapping bias and genotyping errors. To correct for non-genetic confounders, we applied PCA with and without permutation on normalised read counts in log2RPM across all sites and picked the first N components whose explained variances are greater than those from permutation as covariates for Rasqual. Finally, we only considered SNVs found within peaks to determine direct allele specific effect on TF binding of PU.1 and CTCF in neutrophils.

#### Allele specific analysis of PCHI-C

The genotypes of PCHIC donors were obtained from Cambridge Bioresource phase 4 (Illumina core exome chip). We phased the genotype using BEAGLE2 (v2.0.5)^71^ and imputed using Positional Burrows-Wheeler Transform and Haplotype Reference Consortium (release 1.1) as reference panel, via the Sanger imputation service. We then filtered sites for ≥5% minor allele frequency, HWE p-value ≥ 1×10^−6^, ≤5% sample missingness and INFO score > 0. 8. We removed indels and only considered biallelic single nucleotide variants. We used WASP^72^ to remove PCHIC reads that are likely to be biased towards the reference allele. We then mapped deduplicated ChIP-seq reads on each allele of each SNVs using GATK ASEReadCounter with default parameters, base quality ≥2 and mapping quality ≥15. We then filtered for heterozygous SNVs only with ≥ 10 read counts per site and nonzero counts in both alleles. Finally, we only considered heterozygous sites with allele bias of ≤40% or ≥60%, after removing extreme bias of <1% or >100%.

#### Enrichment analysis of tfQTLs and hQTLs in PIRs

Each of these heterozygous SNVs was annotated based on whether they were located in a PIR and whether they were significant tfQTLs (PU.1 and CTCF; p<1×10^−5^) or significant hQTLs (H3K27me3, H3K4me3, H3K27ac; p<1×10^−5^). Fisher’s exact tests were carried out separately for each sample and for each cell type to test for enrichment of tfQTLs and hQTLs that fall into PIRs. Finally, the mean and standard deviation were calculated across all samples for each cell type. In another approach, all samples were combined across both cell types. SNVs were removed if they were not observed in at least two samples, or in one sample and in the two cell types, or if the allelic ratio (REF reads/ALT reads) was not consistent across the samples or cell types. Enrichment was tested for SNVs where at least N samples fell into a PIR and at least N samples carried a significant tfQTL or hQTL for increasing number of samples N (N=1,2,3,4).

#### Enrichment of genome wide association SNPs within ChIP-seq marked regions

To test for significant enrichment of trait associated SNPs within regions of interest, we applied GWAS analysis of regulatory or functional information enrichment with LD (GARFIELD)^48^. H3K27ac and H3K4me1 occupied regions in neutrophils were obtained from^7^. Neutrophil annotations for PU.1, C/EBPβ, H3K4me3 and H3K27me3 were generated as described above. With the exception that H3K27me3, regions were not chunked into 2.5Kb bins. Monocyte annotation are described in Supplementary Figure 3a for PU.1 and C/EBPβ.

#### Colocalisation between diseases and molecular trait

To overlap our QTL results to GWAS catalogue, we calculated the LD information based on our WGS data using plink v1.9^73^. For all the QTLs that either directly mapped to the GWAS variants or in LD (r^2^≥0.8), we considered that the QTL variant overlapped with a GWAS signal. For the cases where we further selected six autoimmune diseases, we took forward the overlapping disease variants with P-value ≤5×10^−8^ in six selected studies are celiac disease [CD]^42^, inflammatory bowel disease [IBD]^43^, including Crohn’s disease [CD] and ulcerative colitis [UC], multiple sclerosis [MS]^44^, Type 1 diabetes [T1D]^45^, and rheumatoid arthritis [RA]^46^. The associations of IBD, CD and UC in the European cohorts were used for this study. We also used Type 2 diabetes^47^ as a negative control. We used a Bayesian colocalization method^49,50^ to elucidate whether the observed overlap between disease and molecular trait may due to a shared genetic effect. The method calculates the posterior probability (PP), versus the null model of no association, for four alternative models: a model where a region or locus contains a single variant associated with either the molecular trait or disease (models 1,2); a model where a single causal variant affects association with both traits (model 3); or a model where two distinct associations exist (model 4). The method derives the PP of each variant in the locus being causal one under different models, and the PP of a given locus is then the integral sum of the PPs of all variants within, with all variants under equal prior probability to be causal. The prior for each model is computed to be one that maximizes the log-likelihood function^50^. We acknowledge the limitations of the model: it assumes one causal variant in the locus; and in the case of high LD between two causal variants the model has limited power to distinguish model 4 from model 3. We also note that colocalization does not imply a causal relationship between molecular trait and diseases, but may be compatible also with the same variant having independent (‘pleiotropic’) effects on molecular traits and disease. We applied colocalization test for each of the 1,003 disease-molecular trait pairs, where the lead SNPs in both traits are in high. r^2^≥0.8. To avoid overlapping 2Mb-wide genetic loci due to features in close proximity (e.g., splicing junctions, genes, histones peaks, CpGs in islands), we tested colocalization per locus, which means that the prior model parameters were estimated using one locus instead of multiple loci and hence the priors may be overestimated.

## Supplementary Tables

**Supplementary Table 1. ChIP-seq data, enrichment and quality control metrics.**

ChIP-seq quality control metrics including number of aligned reads, percent duplicate reads, MACS peaks called. Including metadata for sample identification, antibody and gender.

**Supplementary Table 2. Table of quantitative trait loci results.**

Results from Limix QTL analysis for each feature tested for PU.1, C/EBPβ, CTCF, H3K4me3 and H3K27me3 in neutrophils and the statistical test results for each sentinel SNP.

**Supplementary Table 3. Proximal and distal quantitative trait loci results for sentinel PU.1 SNPs.**

Classification of proximal and distal PU.1 tfQTLs reaching the significance threshold (FDR<0.05).

**Supplementary Table 4. Differentially regulated genes associated with H3K27me3 remodelling QTLs.**

Summary of genes identified as been associated with H3K27me3 remodelling and differential expression during neutrophil terminal differentiation that also contain H3K27me3 hQTLs.

**Supplementary Table 5. PU1 tfQTLs that overlap autoimmune diseases and UK Biobank myeloid traits.**

Summary results from colocalisation analysis for PU.1 tfQTL with selected disease and full blood count traits.

**Supplementary Table 6. Annotation of PU1 tfQTLs.**

PU.1 tfQTL were annotated for significant allele-specific effect (RASQUAL analysis), neutrophil gene expression QTL, baited genes through PCHi-C interactome data and whether QTL summary statistics colocalise with any of the GWAS traits tested.

## Supplementary Figures

**Supplementary Figure 1.**
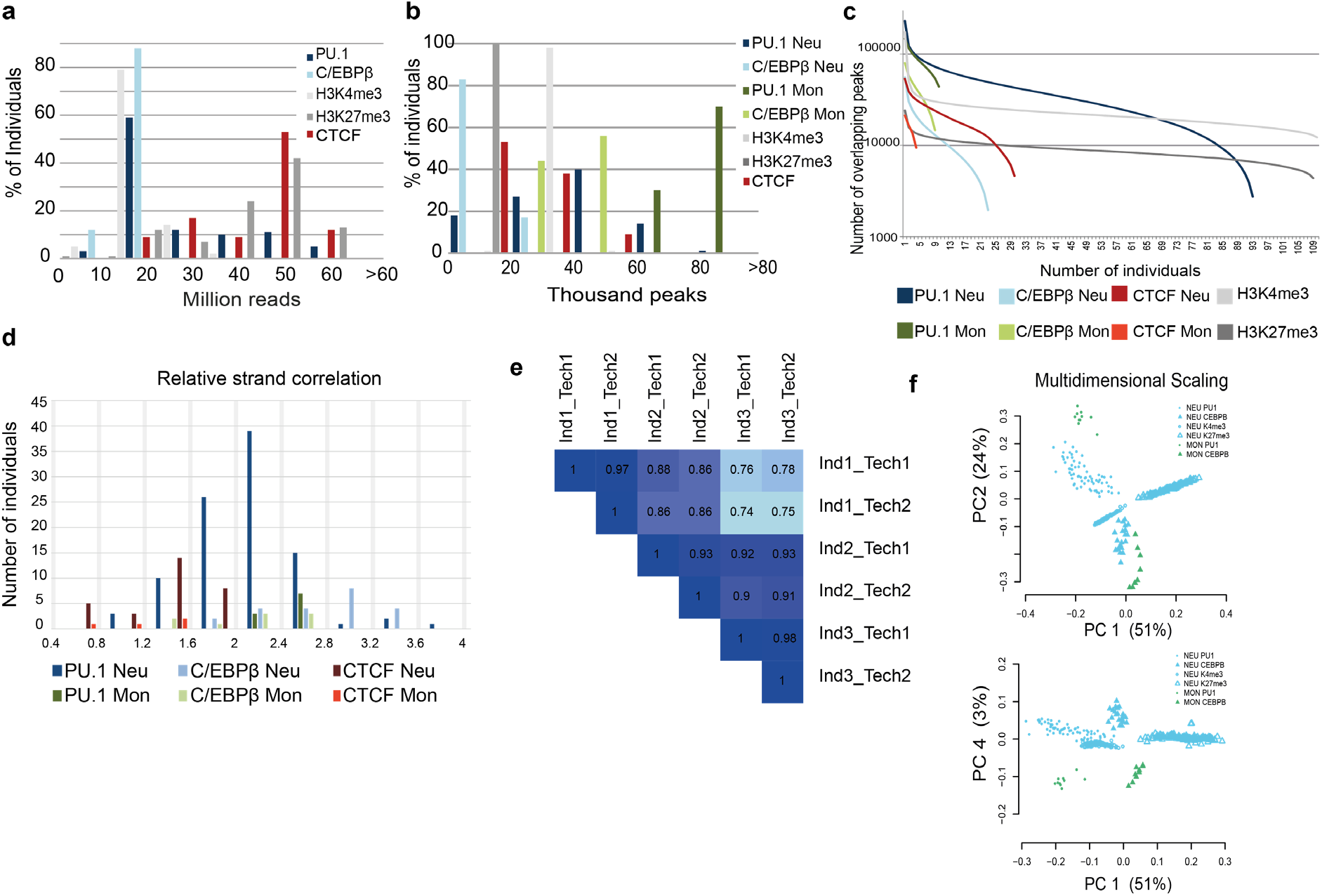
Data QC plots for ChIP-seq data. **a**. Bar plot of bins showing the proportion of individuals with the number of QC passed aligned reads for each factor probed. **b**. Bar plot of bins of number of peaks called for each transcription factor, individual and in the two cell types. Monocytes consistently have more binding locations for both PU.1 and C/EBPβ over neutrophils data sets. **c**. Peak overlap plot split by factor and cell type. Y axis represents the total number of peaks called in all individuals, X axis represents the number of individuals where that peak is called. **d**. Bar plot of bins for the proportion of data sets with a relative strand correlation coloured by factor. **e**. For three individuals, PU.1 profiling was carried out in duplicate as independent technical replicates. Heatmap of pairwise analysis of logRPM signal within a consensus peak set of ~55,000 shared sites, numbers are the Pearson’s correlation between replicates. **f**. Variation in data sets shown by Multidimensional scaling PC1 versus PC2 and PC1 versus PC4.

**Supplementary Figure 2:**
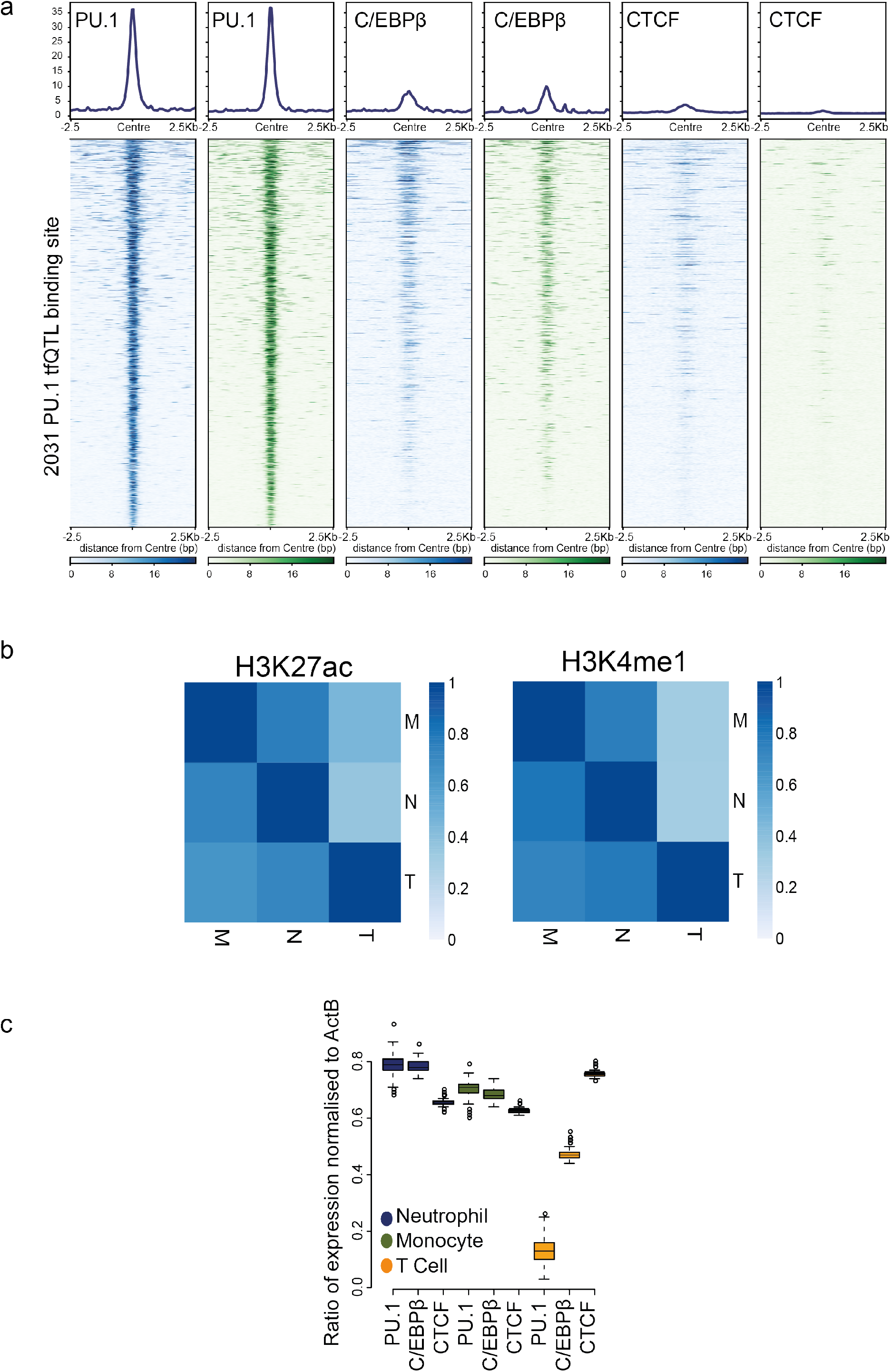
Overlap of PU.1 QTLs with chromatin state. **a**. Sequencing read density heatmap for regions around (+/− 2.5Kb) PU.1 tfQTLs for PU.1, C/EBPβ and CTCF in both neutrophils (blue) and monocytes (green). Percentage of intersecting PU.1 tfQTL peaks which overlap (1bp) with second peak set; PU.1 monocyte 93%, C/EBPβ neutrophil 36%, C/EBPβ monocyte 40%, CTCF neutrophil 11% and CTCF monocyte 5%. **b**. Heatmap of Pi1 statistics of QTL sharing for PU.1 tfQTL across neutrophil, monocyte and T cell types for H3K27ac and H3K4me1 QTLs. **C**. Relative levels of PU.1, C/EBPβ and CTCF gene expression compared to ActB gene from ~200 donors from^7^ across neutrophils, monocytes and CD4 Naïve T cells.

**Supplementary Figure 3:**
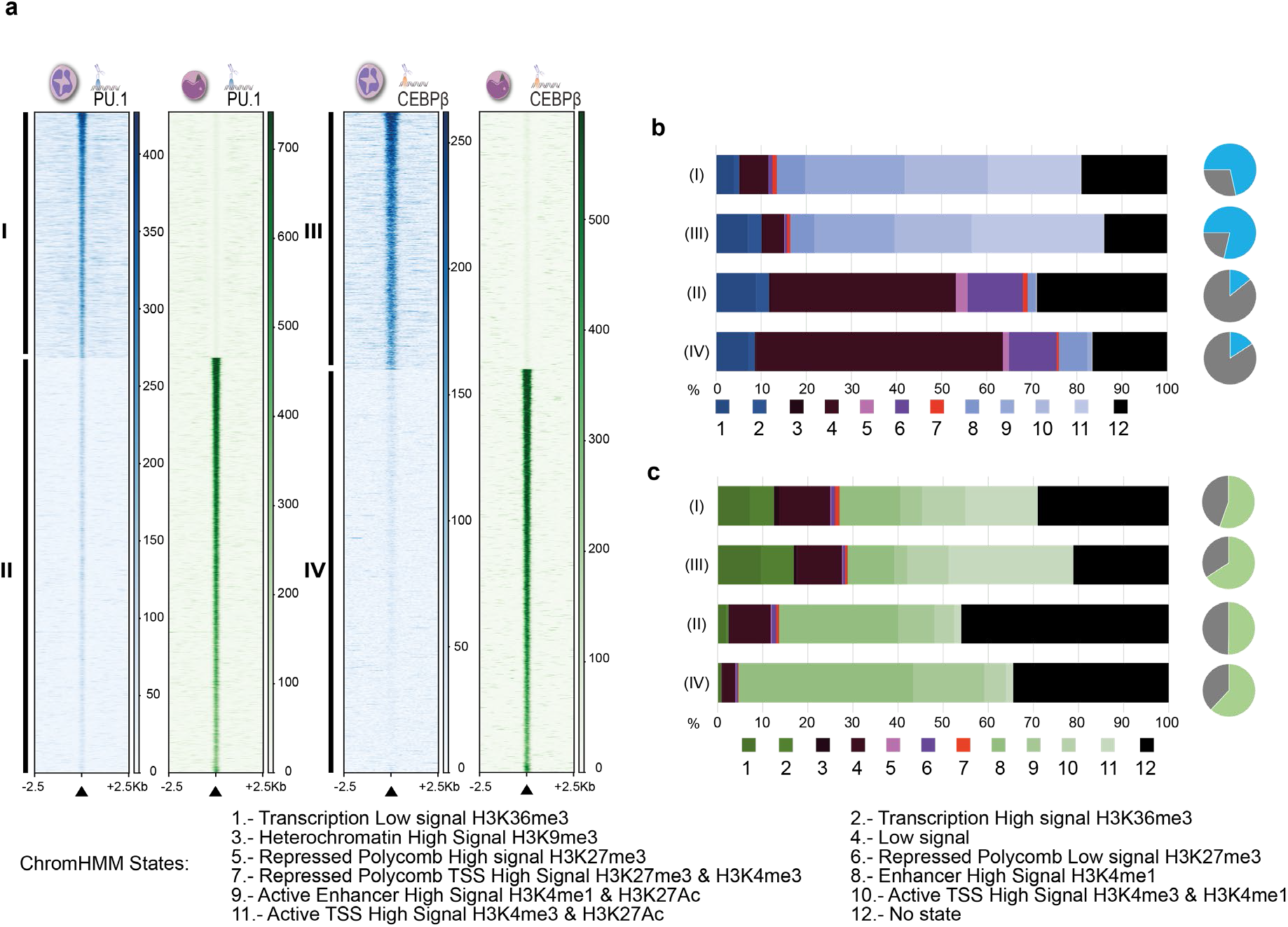
Identification of PU.1 and C/EBPβ differentially bound binding sites. **a**. Pairwise differential binding analysis between Monocytes and Neutrophils for PU.1 and C/EBPβ was performed to identify cell type enriched binding events for the two factors. Read density heatmap +/− 2.5kb from centre of binding site. **b**. Intersection of transcription factor binding sites from A. with chromatin state maps for neutrophils derived using ChromHMM^74^. Pie charts; blue segment is the proportion of TF binding category that fall within regions classed as active in neutrophils. **c**. Intersection of transcription factor binding sites from A. with chromatin state maps for monocytes. Pie charts; green segment is the proportion of TF binding category that fall within regions classed as active in monocytes.

**Supplementary Figure 4.**
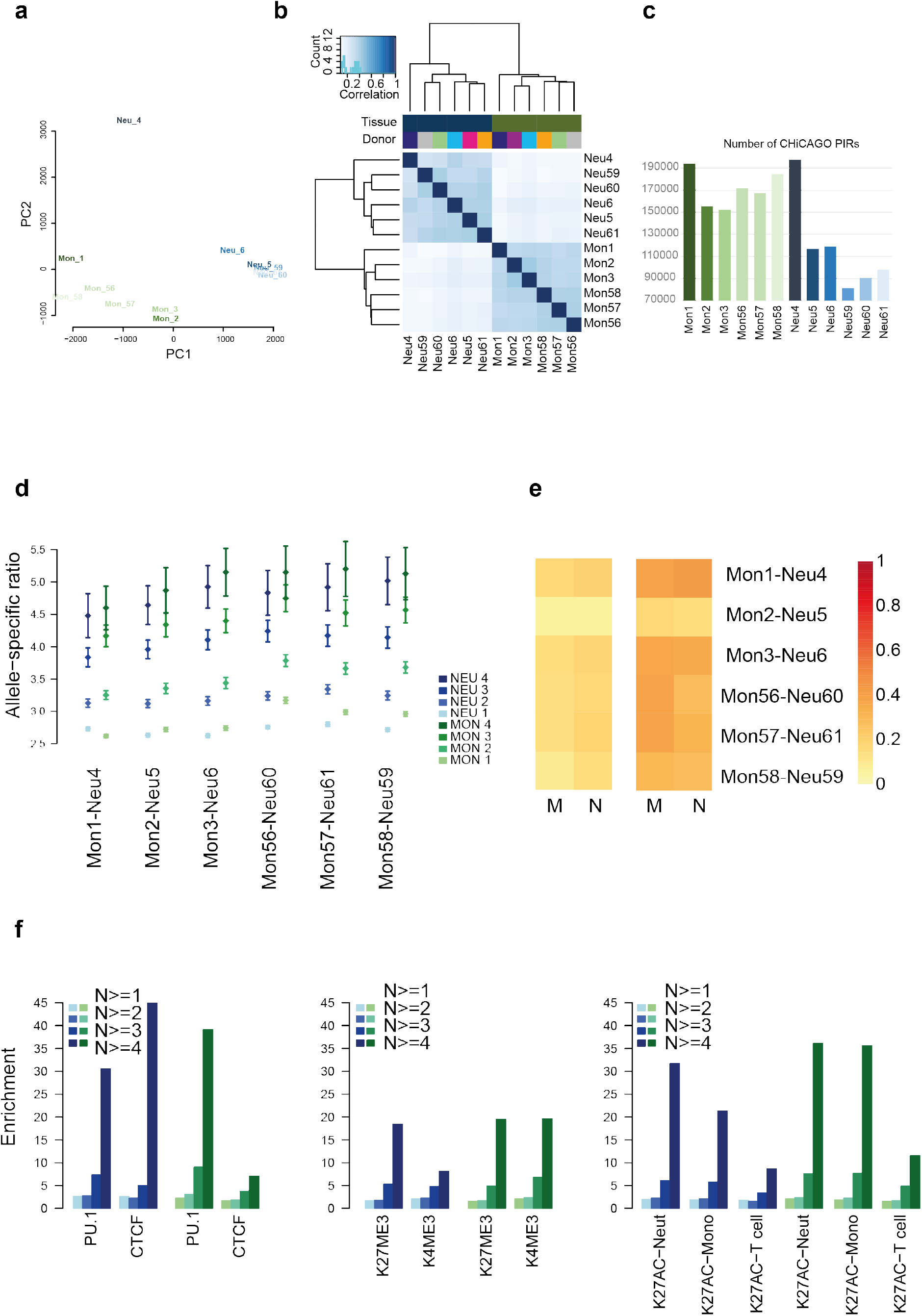
Summary of PCHi-C data in two cell types. **a**. Principal component analysis plot showing first 2 principal components across twelve PCHi-C data sets. **b**. Heat map of Pearson correlations for intersecting (1bp overlap) PIRs (CHiCAGO >5) from twelve data sets. **c**. Bar plot of the number of promoter enhancer connections called with a CHiCAGO score >5 for twelve data sets. **d**. Using the PCHi-C data from seven individuals, we selected 310,233 heterozygous sites in neutrophils (NEU) and 288,385 sites in monocytes (MON) with allele-specific (AS) bias >1.5 or <0.67, though we removed sites with extreme Allele Specific (AS) bias (<0.01 or >100). 89% of sites with AS bias in NEU and 92% in MON had a consistent AS ratio if found in more than one individual. The percentage of sites with consistent AS bias detected in two individuals drops to 16% and 15%, and it drops further to just 3% or to <1% when shared by three or four individuals. The plot shows the mean allele-specific (AS) ratio or bias and the 95% confidence interval by individual in neutrophils and monocytes. The mean AS ratio increases when AS ratios are shared across samples in a consistent manner. In almost all individuals AS ratios are higher in monocytes than in neutrophils. **e**. Left heat map; Percentage of sharing between cell types of SNVs that are located in PIRs. Sharing is shown between monocytes (left) and neutrophils (right) in five matched samples (monocytes mean = 14.4%, neutrophils mean = 18.2%) and one mismatched sample (Mon2 and Neu5), (monocytes mean = 6.0%, neutrophils mean = 6.6%). Right heat map; Percentage of sharing between cell types of SNVs that are significant PU.1 QTLs (p<1×10^−5^). Sharing is shown between monocytes (left) and neutrophils (right) in five matched samples (monocytes mean = 37.0%, neutrophils mean = 33.9%) and one mismatched sample (Mon2 and Neu5), (monocytes mean = 17.0%, neutrophils mean = 15.9%). **f.** Allele-specific SNVs, identified through PCHi-C, were selected if they were observed in at least two samples or cell types, and if their REF/ALT ratio was consistent, i.e. either > 1 or < 1, across all samples. Enrichment of QTLs in PIRs was then calculated for 14,000 SNPs that fulfilled these criteria and that were supported by an increasing number of samples that showed evidence for falling into a PIR and being a significant QTL (N=1,2,3,4). Enrichment increased when increasing the number of supporting samples.

**Supplementary Figure 5.**
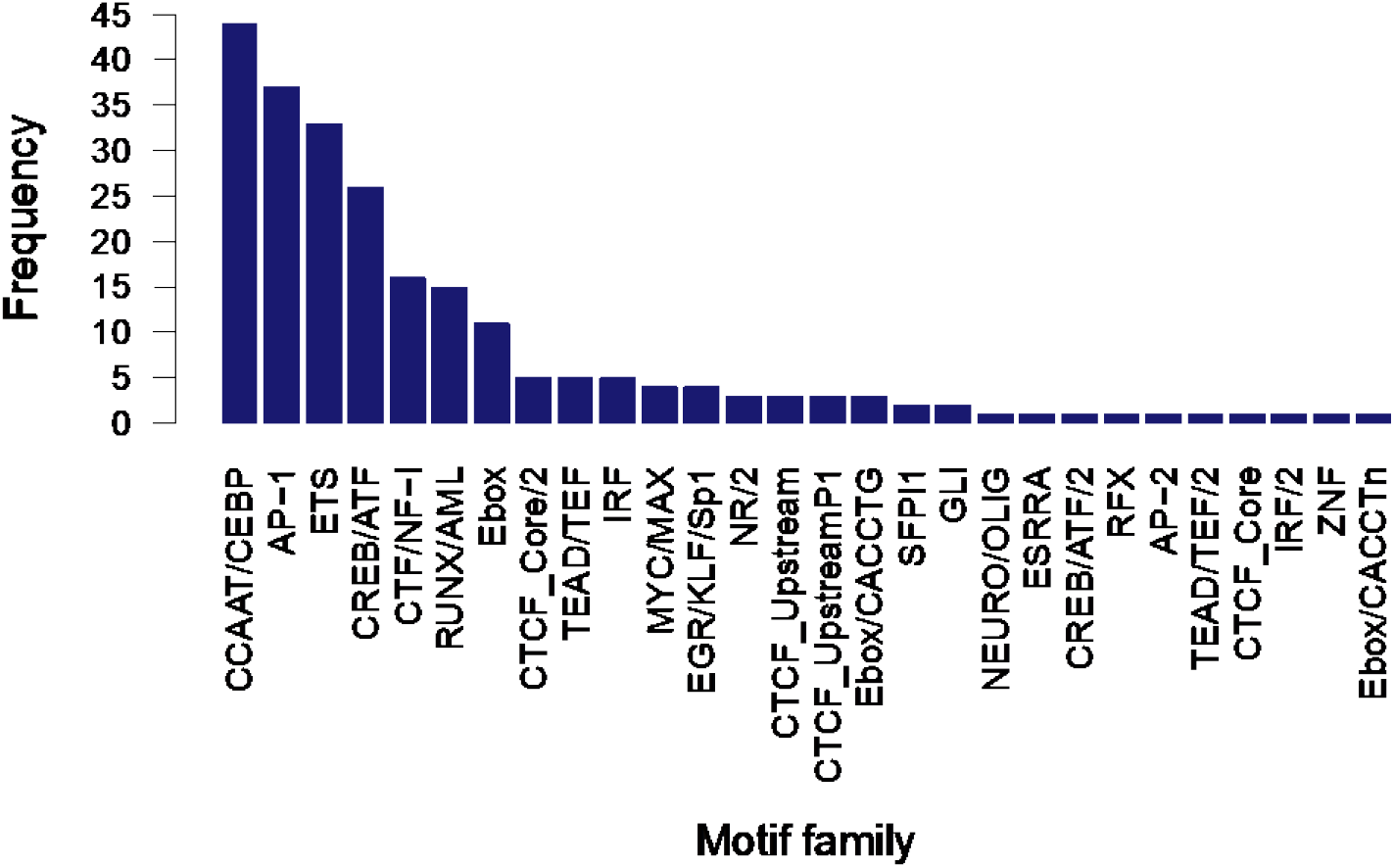
Frequency of motif disruption for transcription factor families at colocalised loci. Bar plot with the frequency of motif families with a predicted transcription factor disruption (CATO score >0.1). The top 6 clusters harbour 79% of the 231 tfQTLs *(i.e.* PU.1 lead tfQTLs and proxies with LD>0.8).

**Supplementary Figure 6.**
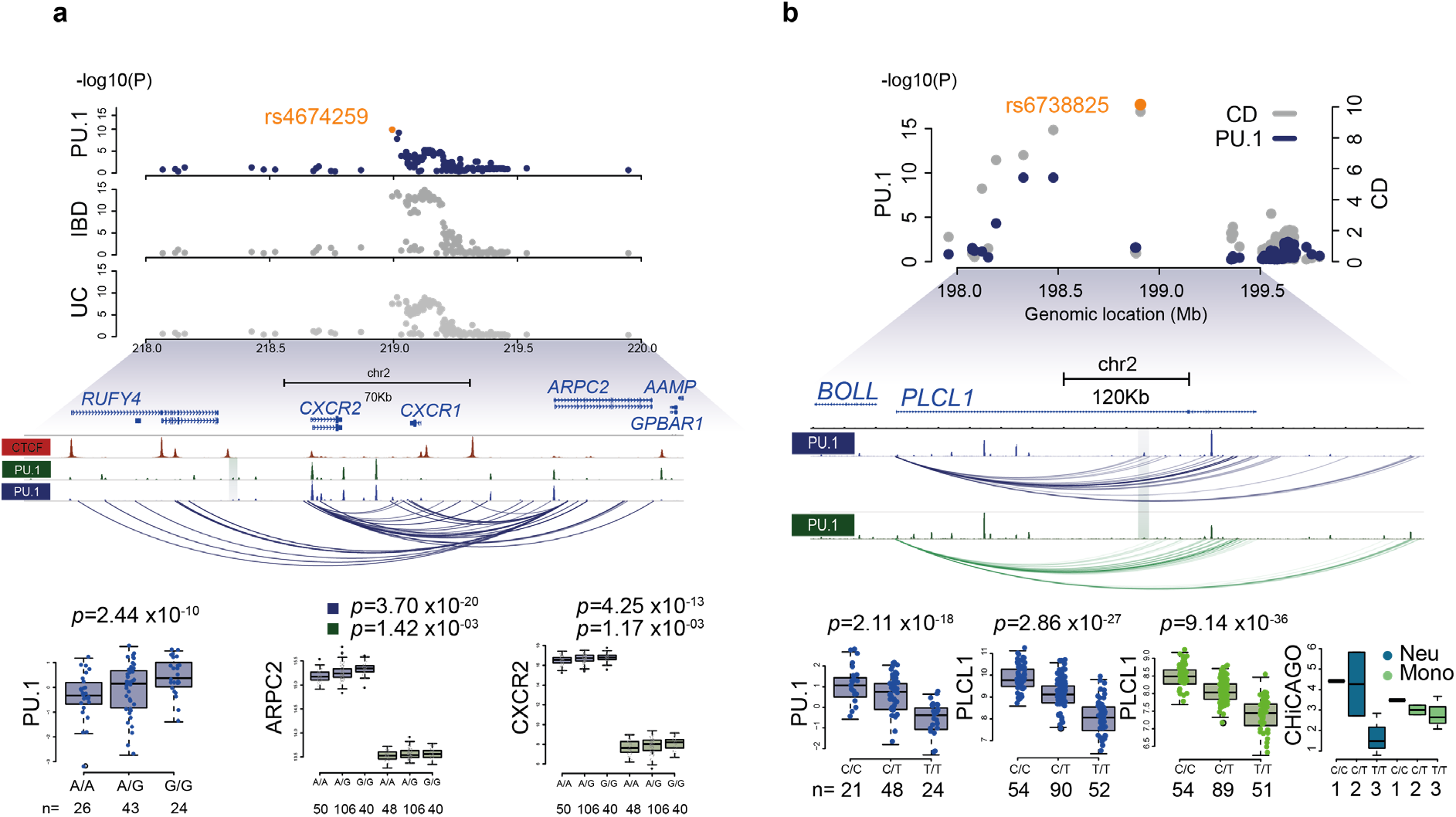
Examples of disease associated loci. **a**. Example of colocalised signal for PU.1 tfQTL, rs13035725 (*p* = 2.44×10^−10^) identifies a risk locus for inflammatory bowel disease and ulcerative colitis. The same SNP is also significantly associated with expression of several genes in neutrophils, including *ARPC2* (*p*=3.70×10^−20^), *CXCR2* (*p*=4.25×10^−13^), *AAMP* (*p*=1.02×10^−8^) and *CXCR1* (*p*=1.23×10^−6^), all of which were weakly or not significant in monocytes. These genes perform highly relevant immune functions for example in neutrophil migration, suggesting disease risk may be mediated through neutrophil biology. The region also displayed high connectivity in the neutrophil PCHi-C data (CHiCAGO score > 5), but depleted of significant interactions in monocytes. In orange location of rs4674259 the most significant shared SNP between tfQTL and GWAS data sets. **b.** Example of colocalised signal for PU.1 tfQTL rs9712275 with Crohn’s disease. Top; Manhattan plot of the-log10(P) values for shared SNPs from PU.1 tfQTL (navy) and Crohn’s disease (grey), position of lead shared SNP rs6738825 highlighted in orange. Middle; genome browser shot of PU.1 binding and promoter interacting regions baited to *PLCL1* gene both neutrophils (navy) and monocytes (green). Lead PU.1 QTL peak is highlighted by shaded area. Bottom; boxplots of signals for molecular traits; PU.1, gene expression and PCHi-C split by donor genotype for rs9712275. The sentinel SNP, rs9712275 (PU.1 QTL *p*=2.11×10^−18^) colocalised with Crohn’s disease and was also associated with active marks, H3K4me3 (*p*=3.91×10^−22^) and H3K27ac (*p*=1.05×10^−19^) as well as the repressive mark H3K27me3 (*p*=1.57×10^−21^). This locus is also a significant eQTL for the phospholipase C, epsilon (PLCL1) gene (*p*=6.11×10^−28^). Interestingly, two associations were also significant in monocytes, H3K27ac (*p*=4.75×10^−15^) and the *PLCL1* eQTL (*p*=3.30×10^−40^). From our allele-specific analysis, we identified that this locus showed evidence of allelic imbalance in PCHi-C interactions for regions interacting with the PLCL1 baited gene. PLCL1 encodes the phospholipase C epsilon or phospholipase C like I (inactive) signalling protein that has been shown to be involved in receptor turnover but also inhibiting integrin activity^75,76^ suggesting a role in the regulation of cell trafficking. Combined these examples, and others, highlight a role for PU.1 mediated regulatory cascade in moderating disease risk.

